# Microstructural underpinnings and macroscale functional implications of temporal lobe connectivity gradients

**DOI:** 10.1101/2020.11.26.400382

**Authors:** Reinder Vos de Wael, Jessica Royer, Shahin Tavakol, Yezhou Wang, Casey Paquola, Oualid Benkarim, Nicole Eichert, Sara Larivière, Bratislav Misic, Jonathan Smallwood, Sofie L. Valk, Boris C. Bernhardt

## Abstract

The temporal lobe is implicated in higher cognitive processes and is one of the regions that underwent substantial reorganization during primate evolution. Its functions are instantiated, in part, by its complex layout of structural connections. This study identified low-dimensional representations of structural connectivity variations in human temporal cortex and explored their microstructural underpinnings and associations to macroscale function. We identified three eigenmodes which described gradients in structural connectivity. These gradients reflected interregional variations in cortical microstructure derived from quantitative MRI and post-mortem histology. Gradient-informed models accurately predicted macroscale measures of temporal lobe function. Gradients aligned closely with established measures of functional reconfiguration and areal expansion between macaques and humans, highlighting the important role evolution has played in shaping temporal lobe function. Our results provide robust evidence for three axes of structural connectivity in human temporal cortex with consistent microstructural underpinnings and contributions to large-scale brain network function.

## Introduction

The human temporal lobe is involved in multiple cognitive, affective, and sensory processes, including memory (Vaz et al., 2019), emotional reactivity (Phelps, 2004), semantic cognition (Ralph et al., 2017), as well as auditory processing (Bonilha et al., 2017). Notably, temporal lobe subregions have been suggested to serve as origins of major organizational and evolutionary axes of the human brain (Goulas et al., 2019; Sanides, 1969), and host structures, such as the middle and superior temporal gyri, that have undergone accelerated functional reconfigurations and areal expansion in primate evolution (Eichert et al., 2020; Mars, Sotiropoulos, et al., 2018; Xu et al., 2020). Collectively, these different aspects suggest that the temporal lobe is a hub implicated in important features of human cognition, and that its study may provide key insights into cortical organization and its phylogenetic basis.

In an attempt to understand the role of the temporal lobe in whole-brain networks, prior studies in non-human animals and human participants have started to delineate the complex connectivity profiles of the temporal lobe. Tract tracing studies in non-human primates have charted short range connections as well as long range tracts of the temporal lobe (Webster et al., 1991), showing distributed connectivity patterns to a diverse territory of cytoarchitectural areas (Beul et al., 2017; Mohedano-Moriano et al., 2015; Morán et al., 1987; Sakata et al., 2019). These findings were complemented by diffusion MRI tractography studies in both non-human primates (Bryant et al., 2020) and humans (Saur et al., 2008), where this non-invasive technique can approximate the course of white matter fiber tracts both *in vivo* and *ex vivo.* Diffusion MRI studies have been performed for all major long range fiber bundles (Howells et al., 2018; Roumazeilles et al., 2020; Smiley & Falchier, 2009), for short range fiber systems (Attar et al., 2020) as well as the superficial white matter (Bodin et al., 2020; Hong, Hyung, et al., 2019; Liu et al., 2016; Oishi et al., 2008).

Beyond the mapping of specific fiber bundles, recent years have seen a shift towards the application of unsupervised approaches that identify and visualize low dimensional eigenmodes in connectivity changes across the cortical mantle – also referred to as connectivity gradients (Huntenburg et al., 2018; Margulies et al., 2016). A gradient perspective describes transitions of brain connectivity in a continuous reference frame, which has been proposed to capture subregional heterogeneity as well as functional multiplicity better than techniques that parcellate cortex into discrete subregions and average potentially variable properties within parcels (Bijsterbosch et al., 2020; Haak & Beckmann, 2020; Mars, Sotiropoulos, et al., 2018). Capitalizing on resting-state functional MRI acquisitions, gradient mapping techniques have previously identified principal axes of neural organization in healthy adults and in non-human primates (Buckner & Margulies, 2019; Guell et al., 2018; Haak et al., 2018; Margulies et al., 2016; Vos de Wael et al., 2018; Xu et al., 2020), and these techniques are increasingly used to study lifespan processes related to aging (Lowe et al., 2019; Bethlehem et al., 2020) and typical as well as atypical childhood development (Ball, Seidlitz, Beare, et al., 2020; Ball, Seidlitz, O’Muircheartaigh, et al., 2020; Hong, Vos de Wael, et al., 2019; Larivière et al., 2019; Park, Hong, et al., 2020). In the temporal lobe, these techniques have previously been applied to structural connectivity information, with the goal of subsequent parcellation (Bajada et al., 2017), to describe the ventral and anterior temporal lobe as a structural connectivity convergence zone (Bajada et al., 2019), and to relate structural connectivity gradients to meta-analytic task activations (Blazquez Freches et al., 2020; Yarkoni et al., 2011).

In the current work, we expanded on these previous findings in three ways:

i. We explored regional associations between structural connectivity gradients in the temporal lobe and measures of intracortical microstructure to assess whether large scale connectivity axes are reflected in the local microcircuits. Prior findings in non-human animals suggest that an area’s cytoarchitectonic properties may be predictive of structural connectivity, but precise associations between both remain underspecified in humans (Barbas, 2015). To fill this gap, our project leveraged both myelin sensitive MRI contrasts as well *as post-mortem* cytoarchitecture analysis based on BigBrain (Amunts et al., 2013).
ii. Structural connectivity is generally assumed to constrain functional connectivity (Deco et al., 2017; Honey et al., 2009; Suárez et al., 2020; Wang et al., 2019). Here we assessed whether structural connectome gradients within the temporal lobe, as a low dimensional representation of structural connectivity, can predict intrinsic functional organization derived from resting-state fMRI acquisitions, both with respect to macroscale functional motifs as well as node-wise estimates of functional connectivity.
iii. Finally, to determine phylogenetic principles of structural connectome organization, we examined whether structural connectivity gradients reflect principal dimensions of primate evolution. To this end, we studied the relationship of gradients with areal expansion and functional reconfigurations from non-human primates to humans (Xu et al., 2020).

Our approach capitalized on multimodal image processing and advanced diffusion tractography analyses. Specifically, we leveraged a high-resolution representation of temporal lobe structural connectivity to resolve subregional heterogeneity in connectivity and multiplicity of potentially overlapping gradients. Our findings were replicated both in a hold-out dataset from the same site, and in a dataset acquired with a different scanner, imaging parameters, and preprocessing pipeline. We have released all codes to replicate the main figures on https://github.com/MICA-MNI/micaopen

## Results

Our main analyses were based on 75 unrelated participants of the Human Connectome Project (HCP) S900 release (Van Essen et al., 2013), a large-scale open-access neuroimaging dataset comprised of healthy young adults *(HCP-Discovery;* n=75; age=29.2+3.6, female=47). We also replicated all findings in a subset of unrelated participants from HCP, *(HCP-Replication;* n=75; age=28.9+4.0, female=44). For each participant, we mapped structural connectivity of each vertex in the gray-white matter interface of the temporal lobe to the entire cortex using high resolution tractography (see *Methods*, for details). To identify structural connectivity gradients, we used nonlinear dimensionality reduction techniques that identify spatial eigenvectors explaining interregional variations in structural connectivity (Coifman & Lafon, 2006). To assess the reproducibility of our findings, we repeated our analyses on the Microstructure Informed Connectomics *(MICs)* dataset, a separate dataset of healthy controls who underwent 3T imaging comparable to the HCP protocol in our center (54 controls, 30.5+7.3 years old, 20 females).

### Multiple gradients of structural connectivity in the temporal lobe

In *HCP-Discovery,* the first three components of temporal cortical gradients collectively explained 67% of variance in temporal lobe structural (**Fig. 1**Error!Reference source not found.**A**). Findings were similar in the left and right hemispheres. We thus present only left hemisphere results in the main figures (for right hemisphere results, see **Supplementary Fig. 1**). Gradient solutions were consistent across the different datasets studied, with absolute correlations between G1-3 of *HCP-Discovery* with G1-3 of *HCP-Replication* and *MICs* exceeding r>0.96.

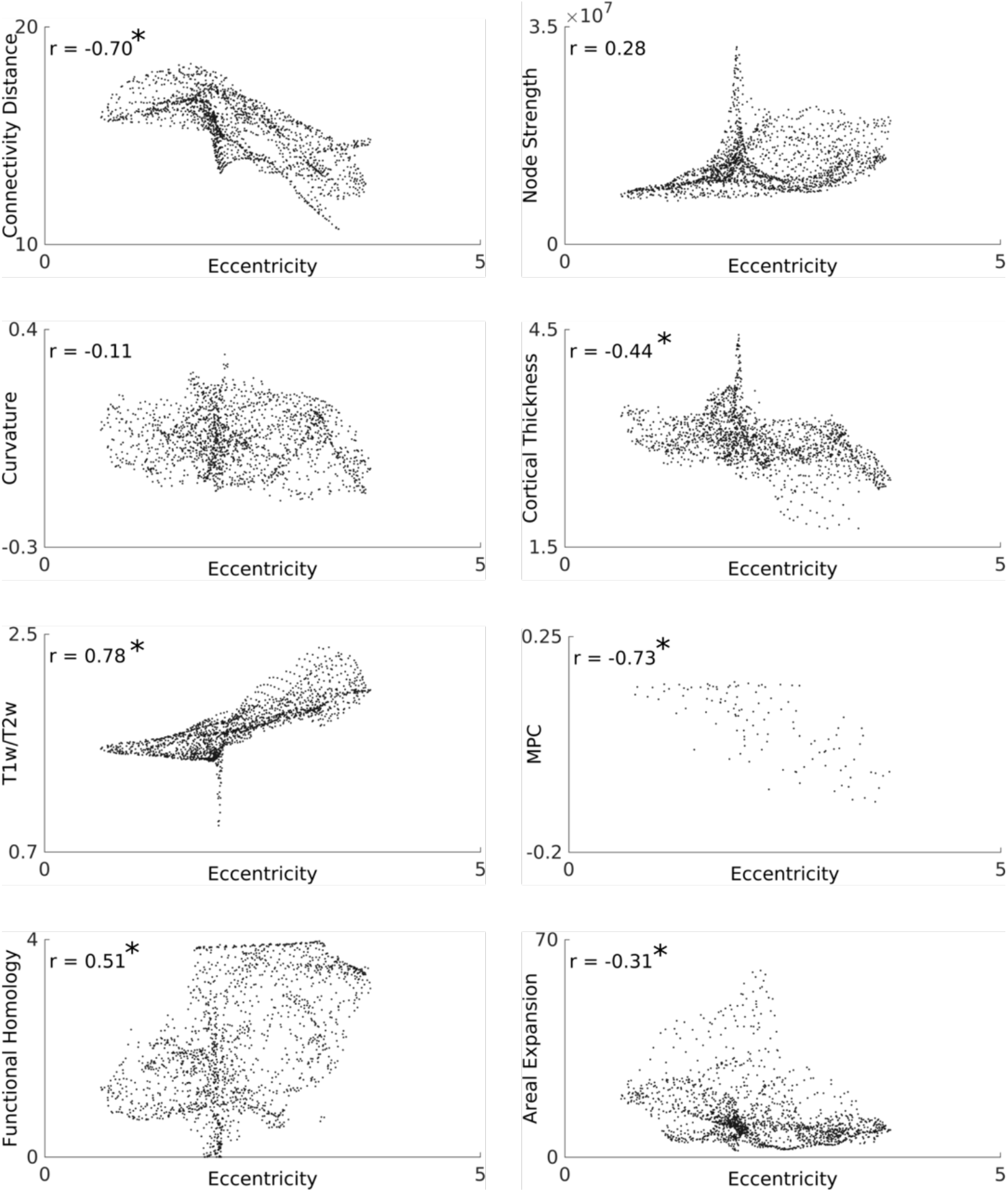

The first structural connectivity gradient (G1) ran between the superior temporal gyrus and the medial temporal lobe (**Fig. 1**Error! Reference source not found.**A**), the second (G2) along the posterior-anterior axis, and the third (G3) from anterolateral to posteromedial. To determine the connectivity patterns represented by each gradient, we mapped the connectivity of the top/bottom 10% of vertices of each gradient and assessed changes in the spatial distribution of connectivity profiles at the anchors of each gradient. G1 connectivity changes differentiated between visual and parietal connectivity, G2 involved changes from temporal pole and insula to visual/parietal cortex and lateral frontal cortex, and G3 described changes from visual/parietal to lateral temporal and frontal (**Supplementary Fig. 2**).

**Fig. 1.**
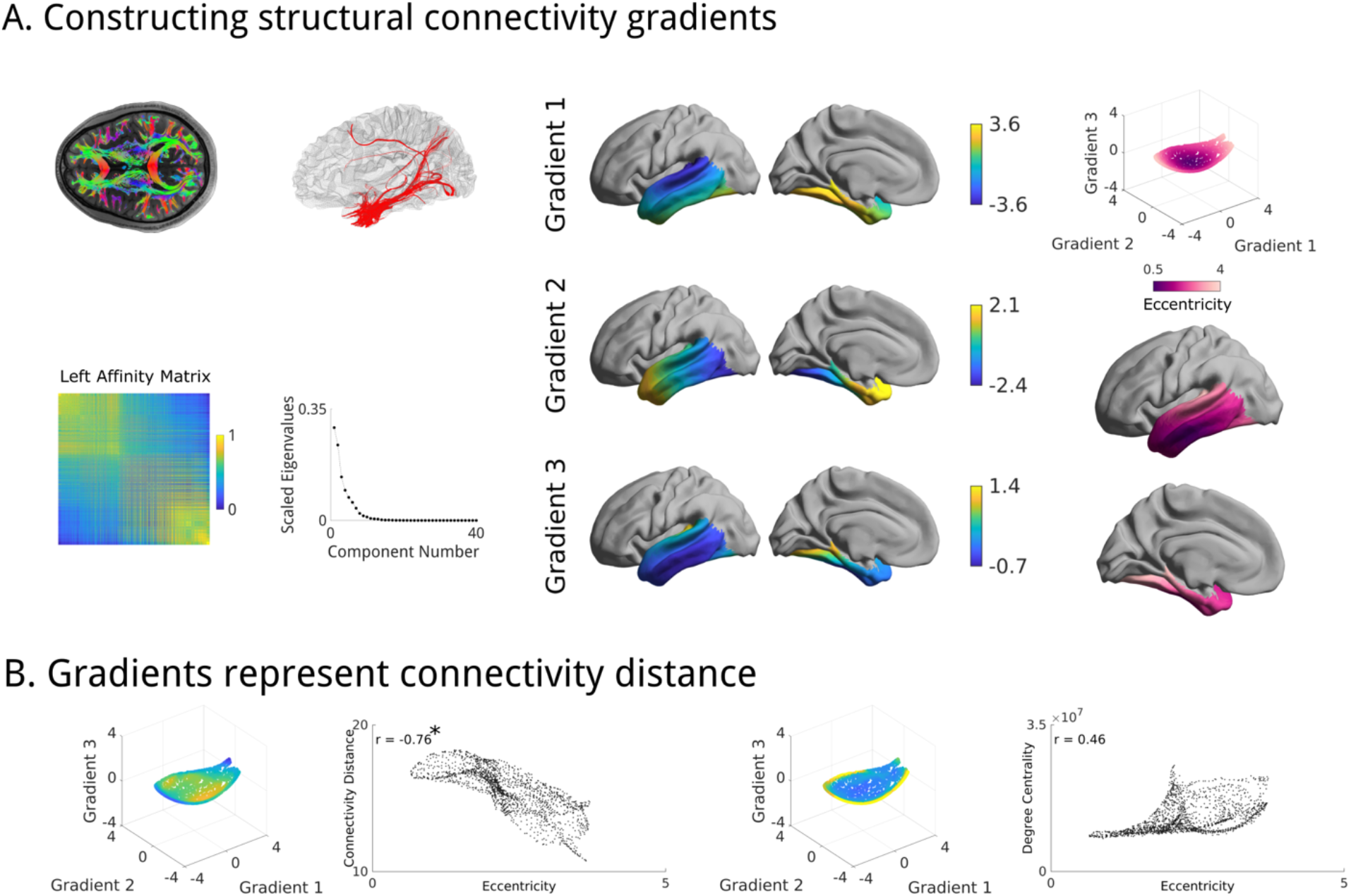
Diffusion embedding of structural connectivity. **(A)**Streamlines were generated throughout the entire brain and systematically mapped to the cortical surface using nearest neighbor interpolation. We computed the affinity matrix of the connectivity matrix using a cosine similarity kernel and constructed gradients of structural connectivity of the temporal lobe to ipsilateral hemisphere with diffusion embedding. The first three gradients of diffusion embedding, sorted by variance explained of each vector, were selected for further analyses. Eccentricity in this manifold space was high in posterior medial temporal lobe and the superior temporal gyrus. **(B)**The relationship of eccentricity with connectivity distance (left) and degree centrality (right). The star denotes a significant correlation.

In order to provide a scalar metric in multivariate gradient space and quantify connectome-level differentiation across the cortical mantle, we calculated an eccentricity measure that captures the distance from the origin in gradient space (Park, Bethlehem, et al., 2020). Low eccentricities were situated in the middle temporal gyrus, while high eccentricity was present in posterior superior temporal and medial temporal regions. To determine the connectivity patterns underlying gradient eccentricity, we performed spatial correlation analyses between eccentricity and topological measures of degree centrality and long-distance connectivity (**Fig. 1B**). Findings were corrected for spatial autocorrelation with Moran Spectral Randomization (Wagner & Dray, 2015) implemented in BrainSpace (Vos de Wael et al., 2020), and adjusted for multiple comparisons using a false discovery rate procedure (Benjamini & Hochberg, 1995). Gradient eccentricity correlated with connectivity distance in both hemispheres (left/right r=−0.76/−0.70, p_moran_<0.002), but not with degree centrality (left/right r=0.46/0.28, p_moran_<0.22). Results replicated in all datasets *i.e.,* gradients were bilaterally associated with connectivity distance (left/right *HCP-Replication:* r=−0.75/0.70, p_moran_<0.002; *MICs:* r=−0.78/0.71, p_moran_<0.01), but not degree centrality *(HCP-Replication:* r=0.44/0.27, p_moran_<0.23, *MICs:* r=0.40/0.39, p_moran_<0.19).

To contextualize the gradients in their cognitive underpinnings, we decoded the structural gradients and eccentricity map using Neurosynth, an ad hoc meta-analysis of previous fMRI studies (Yarkoni et al., 2011) (**Supplementary Fig. 3**). Both G1, G3, and eccentricity represent axes of sensory functions to self-generated cognitive processes (G1: auditory vs memory/navigation terms, G3: cognitive vs auditory terms, eccentricity: cognitive vs perception terms). G2 differentiated stress/affect related terms from visual/word related terms *(e.g.,*“visual” and “word form” vs “stress”, “pain” and “regulation”).

### Microstructural underpinnings

Prior research in non-human animals has shown inter-regional connectivity is predicted by cytoarchitectural similarity (Barbas, 2015), and recent functional MRI work showed correspondence between functional gradients and proxies for intracortical myelin (Huntenburg et al., 2017; Larivière et al., 2019; Paquola et al., 2019; Vos de Wael et al., 2018). Here, we examined the relationship between structural connectivity gradients and *in-vivo* measures of cortical microstructure. Specifically, we tested for associations of gradient eccentricity and intracortical T1w/T2w intensity, a proxy for myelin (Glasser & Essen, 2011), and observed a strong association (**Fig. 2A;** left/right r=0.69/0.78, p_moran_<0.012). Associations to cortical thickness were only moderate (left/right r=−0.43/−0.44, p_moran_<0.024) and those to curvature did not reach statistical significance (left/right r=−0.07/−0.11,It p_moran_<0.65). Similar findings were seen in HCP-Replication (left/right T1w/T2w r=0.68/0.78, p_moran_<0.012; cortical thickness r=−0.45/−0.44, p_moran_<0.020; curvature r=−0.07/0.12, p_moran_<0.637) and in the MICs dataset, which used quantitative T1 relaxometry as a myelin proxy (left/right qT1 r=−0.36/−0.66 p_moran_<0.062; cortical thickness r=−0.33/−0.31, p_moran_<0.08; curvature r=0.01/0.03, p_moran_<0.91).

**Fig. 2.**
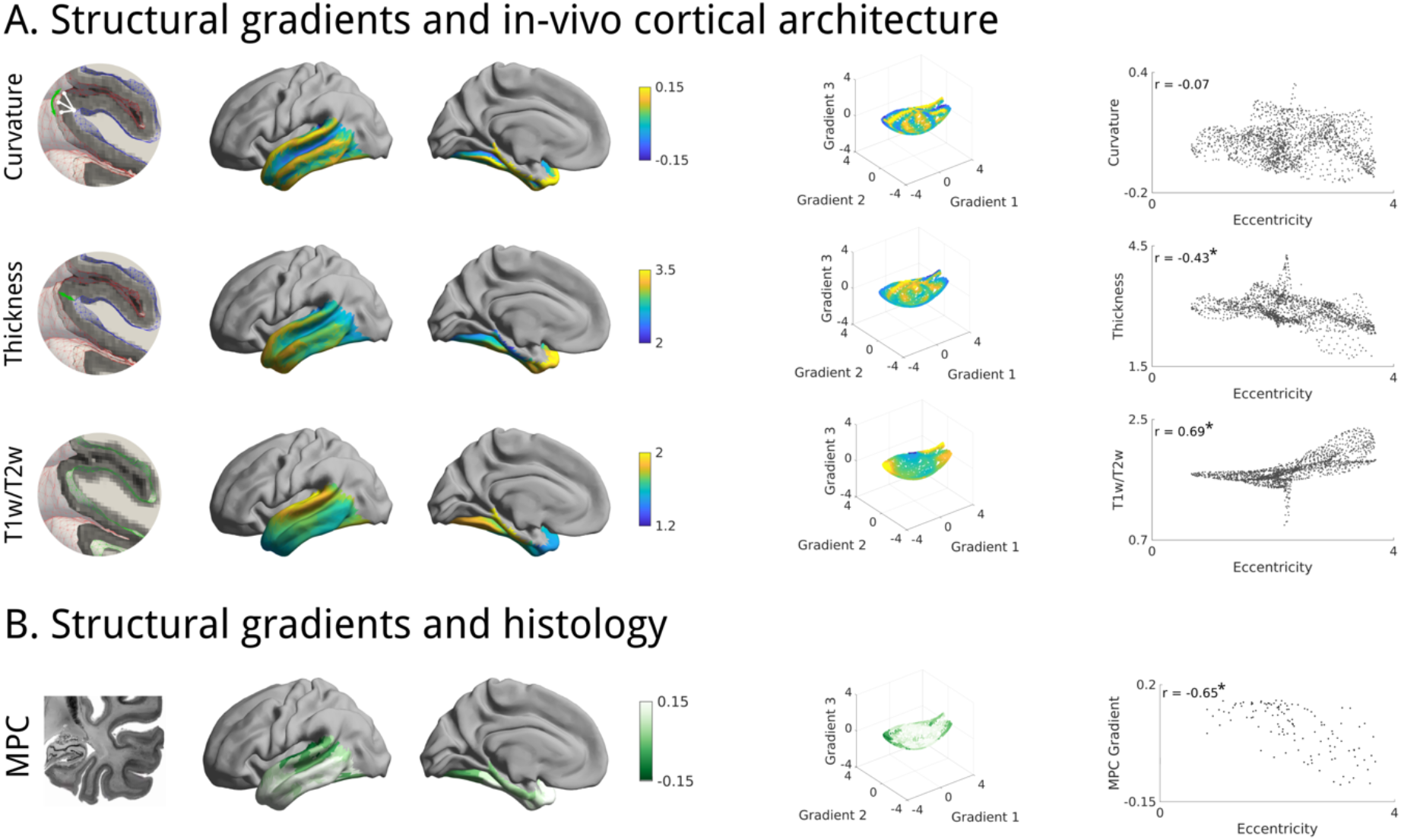
Microstructural basis of the structural gradients. **(A)**We tested for linear relations between eccentricity and curvature, cortical thickness, as well as T1w/T2w intensity. Stars denote significant correlations. **(B)**We tested for an association between microstructural profile covariance derived previously from the BigBrain atlas (Amunts et al., 2013; Paquola et al., 2019), and eccentricity. Eccentricity was projected to the same parcellation scheme as microstructural profile covariance by taking the mean within each parcel.

Next, we evaluated the association between gradients and cortical cytoarchitecture (**Error! Reference source not found.B**), capitalizing on the BigBrain dataset, an ultra-high resolution 3D histological reconstruction of a *post-mortem* human brain (Amunts et al., 2013). We adopted a previously established approach to identify cytoarchitectural gradients (Paquola et al., 2019)and compared the principal cytoarchitectural gradient, which runs from primary sensory to limbic areas, to our *in vivo* structural connectivity gradients. We found strong associations in both hemispheres (left/right r=−0.65/−0.73, p_moran_<0.002). Again, results were replicated in both *HCP-replication* (left/right r=−0.64/−0.73, p_moran_<0.002) and *MICs* (left/right r=−0.60/−0.71, p_moran_<0.028).

### Functional associations

Structural connectivity is ultimately assumed to give rise to functional connectivity (Deco et al., 2017; Honey et al., 2009; Suárez et al., 2020; Wang et al., 2019). As such, we hypothesized that axes of structural connectivity would capture the organization of large-scale functional connectivity. We related the structural connectivity gradients to intrinsic functional community organization, a predominant motif of macroscale neural function (**Fig. A**) (Yeo et al., 2011). Using a 5-fold cross validation, we computed group-level structural gradients for the training and testing group. We derived beta values from the training sets with a group-level multinomial logistic regression and used those to predict the layout of the Yeo-Krienen intrinsic functional communities from the group-level testing set. Predictions were accurate and stable (Cohen’s kappa mean+SD left/right: 0.77+0.01/0.81+0.01). Beta values derived from *HCP-Discovery* gradients also could accurately predict macroscale functional communities from the *HCP-Replication* (Cohen’s kappa left/right: 0.77/ 0.82) as well as the *MICs* (Cohen’s kappa left/right: 0.69/0.70).

To further assess the capacity of gradient features to predict regional functional connectivity, we leveraged decision tree regression with Euclidean distances between vertices in gradient space as predictors and edgewise functional connectivity within the temporal lobe as outcome variable. In a 5-fold cross validation trained on group-level folds of *HCP-Discovery,* gradient space distances were predictive of functional connectivity of held-out subjects at the single subject level (**Fig. 3B**; mean+SD left/right: r=0.50+0.04/0.46+0.05). A decision tree regression trained on the entire *HCP-Discovery* dataset accurately predicted single subject functional connectivity in both *HCP-Replication* (Fig. 3B; left/right r=0.49+0.05/0.43+0.05) and *MICs* (left/right r=0.50+0.03/ r=0.46+0.04). In all datasets prediction quality was excellent in the lateral temporal lobe but less favorable in the medial temporal lobe (**Fig. 3C**), possibly due to lower signal-to-noise ratio in this region.

**Fig. 3.**
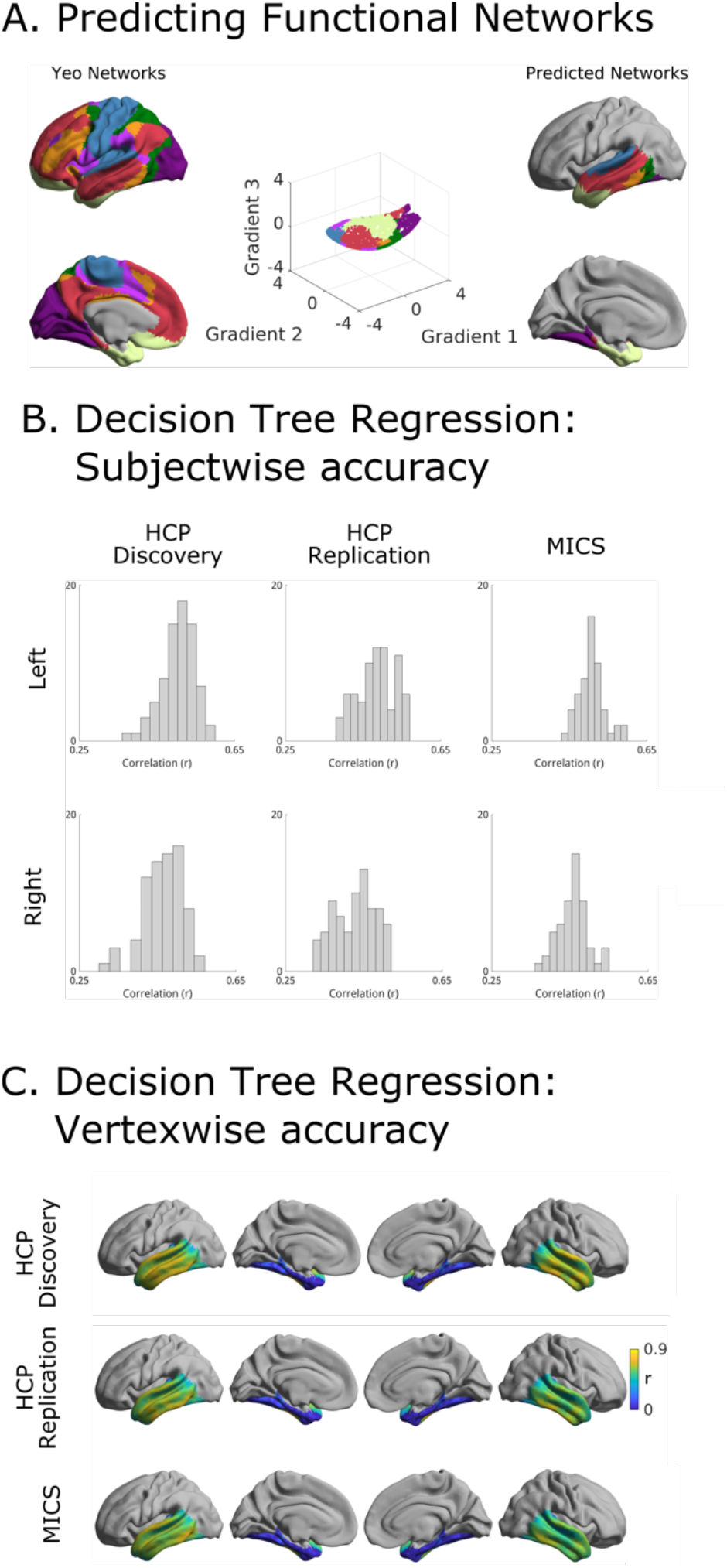
Functional markers of the structural gradients. (A) Based on a canonical network parcellation (Yeo et al., 2011), we attempted to predict the functional networks. Performance was high with a Cohen’s kappa of 0.77+0.01(left) and 0.81+0.01 (right). Predicted networks shown here are the results of one of the five folds. (B) Histograms of the prediction accuracy per subject, as measured by the Pearson correlation between empirical and predicted data, of a decision tree regression estimating functional connectivity from structural gradients. HCP-Discovery predictions were trained with a 5-fold crossvalidation, the predictions of the other datasets were trained on HCP-Discovery (C) Pearson’s correlation between the predicted and empirical functional connectivity for every vertex across subjects. Predictions were especially accurate in lateral temporal regions, and less robust in the medial temporal lobe.

### Evolutionary associations

Last, we assessed whether eccentricity also relates to measures of phylogenetic changes in the temporal lobe (**Fig. 4)**. We found associations to previously established indices of functional homology (left/right: r=0.50/0.51, p_moran_<0.04), a measure for similarity of functional organization between human and macaque, and areal expansion (left/right: r=−0.52/−0.31, p_moran_<0.04), a measure for the surface areas increase of human cortex relative to homologue regions in macaques (Xu et al., 2020). These results replicated both in *HCP-Replication* (functional homology; left/right: r=0.50/0.50, p_moran_<0.04; areal expansion; left/right: r=−0.52/−0.31, p_moran_<0.04) as well as in *MICs* (functional homology; left/right: r=0.48/0.54, p_moran_<0.04; areal expansion; left/right: r=−0.54/−0.40, p_moran_<0.03).

**Fig 4.**
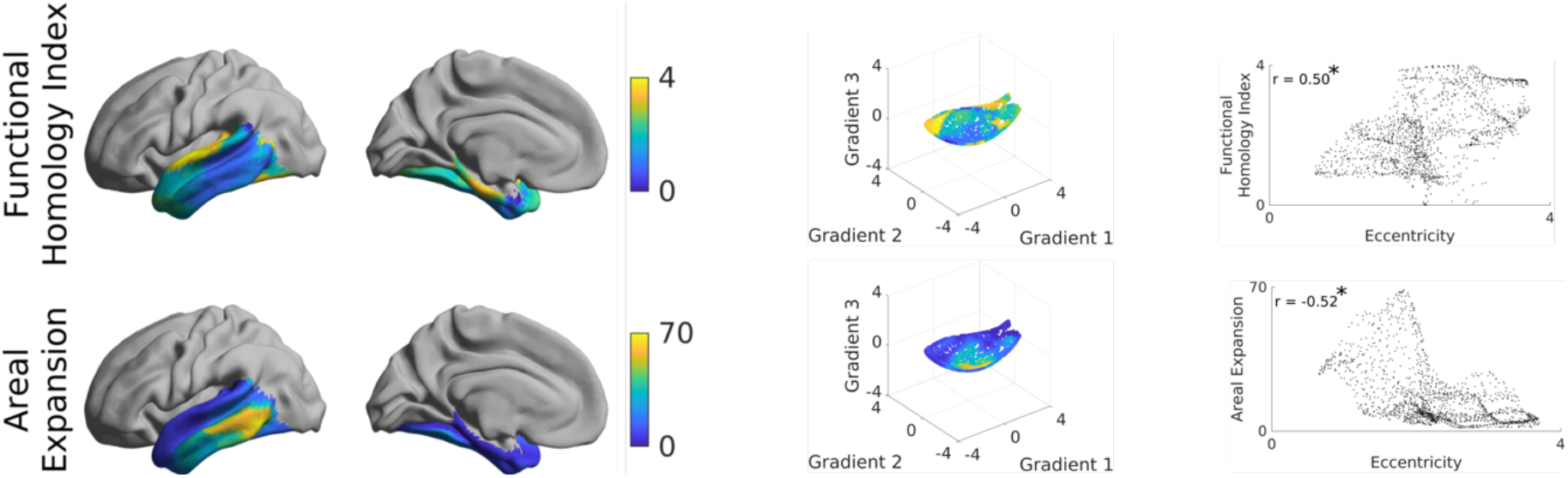
Relationship between the structural gradients and phylogenetic markers. Both functional homology and areal expansion are significantly associated with eccentricity of the first three structural gradients.

## Discussion

The temporal lobe hosts a diverse array of functional processes implicated in sensory processing, memory, and language abilities, and is among the macroscopic structures most frequently compromised in neurological and neuropsychiatric disorders, including Alzheimer’s disease (Braak & Braak, 1991) and drug-resistant epilepsy (Dam, 1982). To provide a comprehensive account of its substructural organization in humans, our study harnessed manifold learning operating on high resolution diffusion MRI tractography data of the temporal lobe to identify separate, yet partially overlapping axes of its structural connectome embedding. These axes were found to relate to MRI-based measures of intracortical myelination as well as *post-mortem* histology, supporting potential microcircuit underpinnings of these spatial trends in structural connectivity variations. Supervised learning experiments indicated that structural gradients can serve as sensitive, low-dimensional predictors of the functional organization of the temporal lobe. Notably, structural connectivity gradients also related to established markers of functional reconfiguration and areal expansion between humans and non-human primates, supporting the potential of connectome gradients in shaping evolutionary changes. Results were reproducible across multiple datasets, indicating generalizability. Collectively, our findings provide robust evidence for an association between tissue microstructure, structural connectivity, and functional motifs of the temporal lobe, which suggests their potential to serve as major organizational axis bridging between its microcircuit and macroscale layout.

Diffusion MRI is the only non-invasive method to approximate the course of white matter connections in humans. Based on multi-shell diffusion acquisitions of the *HCP* and *MICs* datasets, we applied constrained spherical deconvolution (Jeurissen et al., 2014) and spherical-deconvolution informed filtering of tractograms (R. E. Smith et al., 2015a) to estimate streamline weights interconnecting cortical areas. These techniques provide biologically meaningful weights of the modelled streamlines (R. E. Smith et al., 2015b), and reduce fiber tracking biases (Yeh et al., 2016) as well as partial volume effects (Jeurissen et al., 2014). By propagating each streamline to cortical surface points, rather than to macroscopic parcels, we were able to resolve fine grained changes in temporal lobe connectivity and thus account for heterogeneity of subregional connectivity. We enhanced this vertex-wise approach with manifold learning techniques that allow for the low-dimensional representation of spatial variations in temporal lobe structural connectivity. Already established by previous neuroimaging studies (Bajada et al., 2019; Blazquez Freches et al., 2020; Haak et al., 2018; Huntenburg et al., 2017; Margulies et al., 2016; Paquola et al., 2019), these techniques are able to model both gradual and overlapping modes of connectivity without reliance on a priori boundaries (Haak & Beckmann, 2020). Recapitulating prior work, we found that the temporal lobe is best described by three gradients (Bajada et al., 2019; Blazquez Freches et al., 2020), spanning medio-lateral (G1), anterior-posterior (G2) and anterolateral-posteromedial (G3) axes. Although there have been several reports of asymmetry of the white matter tracts of the temporal lobe, such as greater fiber density and tract volume in the left arcuate fasciculus than the right [for review see (Ocklenburg et al., 2016)], the symmetry of the structural gradients indicates gross similarity between large scale network embedding of left and right temporal lobes. We then tested for associations with MRI-based measures of curvature, cortical thickness, and intracortical microstructure. In line with our hypotheses and prior work suggesting a close link between internal cortical architecture and structural connectivity (Beul et al., 2017; García-Cabezas et al., 2019; Scannell et al., 1995; Young, 1992), we found strong associations between connectome gradients and MRI proxies of cortical myelin. The relationship with cortical thickness and curvature was weaker, suggesting that our cortical connectivity gradients closely reflect intracortical factors and only to a lesser extent macroscopic morphological variations and/or potential biases from sulco-gyral folding (Schilling et al., 2018). A closer link to microstructure was also suggested by harnessing BigBrain derived cytoarchitecture gradients (Amunts et al., 2013; Paquola et al., 2019). Collectively, these findings highlight the close relationship between microstructure and structural connectivity, supporting the extension of the structural model of connectivity to humans in the temporal lobes (Beul et al., 2017; García-Cabezas et al., 2019; Scannell et al., 1995; Young, 1992).

Many studies found that structural connectivity may predict functional connectivity by assuming that the strength of functional interactions depends, in part, on the density and efficacy of both direct and indirect structural connections (Deco et al., 2017; Honey et al., 2009; Wang et al., 2019). We hypothesized that the structural gradients, despite their low dimensionality, would still accurately describe functional interactions. Supervised learning approaches with cross-validation could show that gradient-informed models predicted the spatial layout of previously described intrinsic functional communities in the human brain (Yeo et al., 2011). At a more local scale, gradients could also predict patterns of inter-regional functional connectivity, even when trained and tested on datasets acquired from different scanners. Overall, our results support that lowdimensional eigenmode representations of structural connectivity may potentially underpin intrinsic functional architecture of the human connectome. Such a conclusion is in line with several prior studies in healthy individuals showing that whole-brain structural connectivity gradients shape dynamic signaling at rest (Park et al., 2021) as well as dynamic brain reconfigurations during tasks (C. Murphy et al., 2019). In the study of brain diseases associated with macroscale dysfunction, connectivity gradients have furthermore been used to contextualize changes in brain network architecture (Larivière et al., 2020; Li et al., 2020), supporting their utility to serve as coordinate systems of macroscale functional interactions in healthy and diseased brains.

Cross-species comparisons provide a potential window into human uniqueness by studying brain reconfigurations between humans and non-human primates (Buckner & Krienen, 2013; Krubitzer, 2007). Whilst a remarkable conservation of macroscale organizational principles between macaques and humans is evident (Glasser et al., 2014; Margulies et al., 2016; Valk et al., 2020), association cortices have specifically undergone a striking expansion in relative surface and potential participation in distributed functional networks (Buckner & Krienen, 2013; Hill et al., 2010; Mars et al., 2017; Mueller et al., 2013; Patel et al., 2015). Here, we showed that structural connectivity gradients spatially align with the pattern of evolutionarily diverging brain areas and areal expansion, an index for relative areal size differences across species (Xu et al., 2020). Areas near the center of the structural manifold were less functionally homologous and have undergone more expansion relative to macaques. This may indicate that evolutionary changes have preferentially occurred along particular fiber tracts including, for example, the arcuate fasciculus which has undergone critical anatomical modifications between non-human and human primates (Ardesch et al., 2019; Eichert et al., 2019, 2020; Mars, Eichert, et al., 2018; Mars et al., 2013; Rilling et al., 2008). When taken together with the cognitive terms from the Neurosynth metaanalysis, these results indicate that phylogenetic differences in the temporal lobe are primarily situated along those tracts associated with self-generated cognitive processes.

Theoretical accounts, empirical findings, and gradual changes in research culture have increased the scientific value of replications in neuroscience (Ioannidis, 2005; Moonesinghe et al., 2007; Open Science Collaboration, 2015). Here, we replicated our findings in two datasets: 1) a set of unrelated young adults derived from the same dataset as the discovery set *(HCP-Replication)* as well as 2) a separate dataset acquired at the Montreal Neurological Institute (*MICs*). Even after stringent corrections for both spatial autocorrelation (Wagner & Dray, 2015) and multiple comparisons (Benjamini & Hochberg, 1995), most findings held across all datasets indicating good reproducibility. We have released all utilized feature data and associated analysis scripts (https://github.com/MICA-MNI/micaopen), for independent verification of our results and followup analysis. We hope that these data and associated findings continue to pave the way into studying the important relationship between the microstructure, connectivity, and evolutionary development of the temporal lobe.

## Methods

### Participants

We selected 150 unrelated participants from the Human Connectome Project dataset for whom all resting-state, diffusion weighted imaging and structural scans were available and completed in full (Van Essen et al., 2013). These participants were split into *HCP-Discovery* (n=75; age=29.2+3.6, female=47) and *HCP-Replication* (n=75; age=28.9+4.0, female=44) datasets. For the *MICs* dataset, all data were collected in a single testing session per participant between April 2018 and March 2020. Participants (n=54; 30.5+7.3, female=20) all provided informed consent. Participants reported no history of neurological illness. The study was approved by the Ethics Committee of the Montreal Neurological Institute and Hospital.

### Image Acquisition

*a) HCP.* Images were acquired on the customized Siemens 3T “Connectome Skyra”. Two T1w images were acquired with a 3D MPRAGE sequence with the following parameters: TR=2400 ms, TE=2.14 ms, flip angle=8 deg, FOV=224×224mm^2^, voxel size=0.7mm isotropic. Two T2w images were acquired with identical parameters except for the following: TR=3200ms, TE=565ms, variable flip angle. Four resting-state fMRI images were acquired with a gradient-echo echo-planar imaging (EPI) sequence (TR=720ms, TE=33.1ms, flip angle=52 deg, FOV=208×180 mm, 2 mm isotropic voxels, and 1200 volumes per run). Diffusion images were acquired with a spin-echo EPI sequence (TR=5520 ms, TE=89.5 ms, flip angle=78 deg, FOV=210×180 mm, 1.25mm isotropic voxels, b-values 1000, 2000, and 3000 s/mm^2^, 90 diffusion weighting directions). Six diffusion image scans were acquired each lasting 9 minutes and 50 seconds. Half the runs were acquired with left-to-right phase encoding and the other half with right-to-left.

*b) MICs.* Images were acquired on a 3T Siemens Magnetom Prisma-Fit equipped with a 64-channel head coil. Two T1w scans were acquired with a 3D-MPRAGE sequence (0.8mm isotropic voxels, matrix=320×320, 224 sagittal slices, TR=2300ms, TE=3.14ms, TI=900ms, flip angle=9°, iPAT=2). Quantitative T1 (qT1) relaxometry data was acquired using a 3D-MP2RAGE sequence (0.8mm isotropic voxels, 240 sagittal slices, TR=5000ms, TE=2.9ms, TI 1=940ms, T1 2=2830ms, flip angle 1=4°, flip angle 2=5°, iPAT=3, bandwidth=270 Hz/px, echo spacing=7.2ms, partial Fourier=6/8). We combined two inversion images for qT1 mapping to minimise sensitivity to B1 inhomogeneities and optimize intra- and inter-subject reliability (Haast et al., 2016; Marques et al., 2010). DWI images were obtain with spin-echo EPI, including three shells with b-values 300, 700, and 2000s/mm^2^ and 10, 40, and 90 diffusion weighting directions per shell, respectively (TR=3500ms, TE=64.40ms, 1.6mm isotropic voxels, flip angle=90°, refocusing flip angle=180°, FOV=224×224 mm^2^, slice thickness=1.6mm, multiband factor=3, echo spacing=0.76ms, number of b0 images=3). One 7 min rs-fMRI scan was acquired using multiband accelerated 2D-BOLD EPI (TR=600ms, TE=30ms, 3mm isotropic voxels, flip angle=52°, FOV=240×240mm^2^, slice thickness=3mm, multiband factor=6, echo spacing=0.54 ms). Participants were instructed to keep their eyes open, look at a fixation cross, and not fall asleep. Two spin-echo images with reverse phase encoding were acquired for distortion correction of the rsfMRI scans (phase encoding=AP/PA, 3mm isotropic voxels, FOV=240×240mm^2^, slice thickness=3mm, TR=4029ms, TE=48ms, flip angle=90°, echo spacing=0.54ms, bandwidth=2084Hz/Px).

### Structural Preprocessing

*a) HCP.* Structural images underwent standard HCP preprocessing (Glasser et al., 2013). In short, T1w images were corrected for gradient nonlinearity. Repeated scans were co-registered and averaged. After brain extraction and readout distortion correction, T1w and T2w images were coregistered using rigid body transformations. Non-uniformity correction based on the T1w and T2w contrasts was applied. Preprocessed images were nonlinearly registered to MNI152 space. Cortical surfaces were extracted using FreeSurfer 5.3.0-HCP (Dale et al., 1999; Fischl, Sereno, & Dale, 1999; Fischl, Sereno, Tootell, et al., 1999), with minor modifications to incorporate information from both T1w and T2w scans. Cortical surfaces were aligned using MSMAll to the Conte69 template (Robinson et al., 2014).

*b) MICs.* Data were preprocessed with a Freesurfer 6.0 recon_all pipeline. Both native T1w scans were provided as input and combined through this pipeline. Manual corrections of the pial and white matter surfaces were performed for all subjects. Curvature and cortical thickness estimates were generated by the recon_all pipeline. To acquire tissue segmentations for anatomically constrained tractography, the same images underwent a separate pipeline which included linear alignment of both T1w scans, non-uniformity correction, and intensity normalization. Corrected images were segmented into tissue types using MRtrix3’s 5ttgen (R. E. Smith et al., 2012). qT1 images were linearly aligned to the cortical surface using boundary based registration (Greve & Fischl, 2009). qT1 values were interpolated to the surface by taking the average of seven trilinear interpolations evenly interspersed between the 20^th^ and 80^th^ percentile distances from the pial to white matter surfaces using Freesurfer’s *mri_vol2surf* command.

### Resting-State Preprocessing

*a) HCP.* Data underwent standard HCP preprocessing (Glasser et al., 2013). In short, the timeseries were corrected for gradient non-linearity and head-motion. The R-L/L-R scan pairs we used to correct for geometric distortions. Resulting images were warped to the structural image using rigid body and boundary-based registrations. This warp was concatenated with the warp from T1w image space to MNI152 space to transform functional images to MNI152 space. Further processing removed the bias field, removed the skull, and normalized whole brain intensity. A high pass filter (>2000s FWHM) was used to correct for scanner drift, and additional noise was removed using ICA-FIX (Salimi-Khorshidi et al., 2014).

*b) MICs.* The first five volumes were discarded to allow for magnetic field saturation. Images were then reoriented, and motion and distortion corrected. Nuisance variable signal was removed using an ICA-FIX classifier trained on this dataset and subsequent spike regression (Salimi-Khorshidi et al., 2014). Further tissue-specific signal regression was not performed (K. Murphy & Fox, 2017; Vos de Wael et al., 2017). A warp to the Freesurfer T1w image was computed by averaging volumetric timeseries across the time dimension and registering this image using boundary-based registration. Timeseries were sampled to the surface by taking the average of seven trilinear interpolations evenly interspersed between the 20^th^ and 80^th^ percentile distances from the pial to white matter surfaces.

### Diffusion Preprocessing

a) *HCP.* Images underwent standard HCP preprocessing (Glasser et al., 2013). In short, image intensity was normalized across scans. The topup and eddy algorithms were used to correct for EPI distortions, eddy currents, and motion. A gradient nonlinearity correction was performed, and the deviation of the b-values and b-vectors was computed. Mean b0 images were registered to the T1w image with boundary-based registration (Greve & Fischl, 2009), and this registration was used to transform DWI images to T1w space. The brain was masked based on a Freesurfer segmentation.

b) *MICs.* Data were preprocessed and denoised with MRTrix3’s dwipreproc, which is based on FSL’s eddy correction and topup, and dwidenoise (Andersson et al., 2003; S. M. Smith et al., 2004; Tournier et al., 2012). Freesurfer segmentations were registered to the subject’s DWI space using boundary-based registration (Greve & Fischl, 2009).

### High resolution diffusion tractography and gradient mapping

Tractography was performed identically for the HCP and MICs dataset with MrTrix3 (Tournier et al., 2012). Response functions for each tissue type were estimated using the dhollander algorithm (Dhollander et al., 2016). Fiber orientation distributions were modelled with multi-shell multitissue spherical deconvolution (Jeurissen et al., 2014) and subsequently underwent multi-tissue informed log-domain intensity normalization. The structural T1w image was segmented into five tissue types (R. E. Smith et al., 2012). Anatomically constrained tractography was performed systematically for each temporal lobe voxel in the gray-white matter interface by generating streamlines using second order integration over fiber orientation distributions (Tournier et al., 2010). Streamlines were seeded dynamically from the white matter using the SIFT model (R. E. Smith et al., 2015a). Streamline generation was aborted when 40 million streamlines had been accepted. Each streamline was assigned a weight by optimizing a cross-section multiplier derived with the SIFT2 algorithm (R. E. Smith et al., 2015a). Streamline termini were assigned to their nearest vertex on the surface of the gray-white matter interface. Streamlines of which either terminus was further than 3mm from its nearest vertex were discarded. Connectomes were smoothed on the surface using a 20mm full width at half maximum Gaussian smoothing kernel.

To describe the largest axes of variance in connectivity we used diffusion map embedding (Coifman & Lafon, 2006), a non-linear dimensionality reduction techniques technique used previously to identify neocortical, hippocampal, and cerebellar functional gradients (Guell et al., 2018; Margulies et al., 2016; Vos de Wael et al., 2018). Gradients were computed and aligned using the BrainSpace toolbox (https://github.com/MICA-MNI/BrainSpace) (Vos de Wael et al., 2020), with the following settings: sparsity thresholding at the 75^th^ percentile, a cosine affinity kernel, diffusion map embedding dimensionality reduction with α=0.5, and automated diffusion time estimation. Gradient computations were performed separately on left and right temporal lobes. Interhemispheric connections were not included in the gradient computation. Left and right gradients were aligned with Procrustes alignment (Langs et al., 2015) as implemented in BrainSpace. Eccentricity was computed from the aligned gradients by computing the Euclidean distance to the origin of the manifold space spanned by the first three gradients.

### Statistical testing

Testing for linear associations between cortical markers and gradients likely leads to biased test statistics due to the spatial autocorrelation of MRI data violating the independence of observations assumption (Alexander-Bloch et al., 2018). Instead, for each statistical test we generated random datasets with comparable spatial properties. Specifically, we generated random datasets with equivalent spatial autocorrelation as the response variable using Moran spectral randomization with the singleton procedure (Wagner & Dray, 2015) as implemented in BrainSpace (Vos de Wael et al., 2020). All linear models were fitted for the original data as well as 1000 corresponding simulated datasets. Presented p-values were derived from the percentile rank of the true F-statistic in the distribution of F-statistics in the simulated data. Multiple comparison were corrected for false discovery rate with the Benjamini-Hochberg procedure (Benjamini & Hochberg, 1995).

### Tractography analyses

Connectivity distance, a measure that characterizes the relationship between physical distance and connectivity (Larivière et al., 2020), was computed by thresholding the structural connectivity matrix at the 80^th^ percentile, multiplying each connection by the geodesic distance between their nodes, and averaging all connections for each node. Degree centrality was computed as the column-wise sum of the connectivity matrix. Statistical significance of the association between both degree centrality as well as connectivity distance with the gradients was assessed with Moran spectral randomization (Wagner & Dray, 2015).

### BigBrain Gradient

To assess histological properties of the brain we used BigBrain, an ultrahigh resolution atlas of a single post-mortem brain stained for cell bodies (Amunts et al., 2013). Gradients of microstructural profile covariance were computed as described previously (Paquola et al., 2019). In short, 18 equivolumetric surfaces were constructed between the outer and inner cortical surfaces. To reduce partial volume effects, the inner cortical surface was removed. A linear model implemented in SurfStat (Worsley et al., 2009) was used to correct for anterior-to-posterior increases in intensity values (Amunts et al., 2013). Surface vertices were grouped into 1012 parcels which respected the boundaries of the Desikan-Killiany atlas (Desikan et al., 2006; Hong et al., 2017). A microstructural profile covariance matrix was constructed by computing the Pearson correlation of every pair of vectors whilst controlling for the average whole-cortex intensity profile. Gradients were constructed from this matrix using BrainSpace default parameters (90% sparsity, normalized angle kernel, diffusion map embedding, α=0.5, automated diffusion time estimation). To compare structural connectivity gradients to BigBrain gradients, the structural gradients were parcellated using the same parcellation scheme. Moran spectral randomization (Wagner & Dray, 2015) was used to test for associations between BigBrain gradient 1 and the structural gradients.

### Functional Predictions

Structural gradients were used to predict canonical resting-state networks published previously [(Yeo et al., 2011); https://surfer.nmr.mgh.harvard.edu/fswiki/CorticalParcellation_Yeo2011]. HCP-Discovery was split into five folds of 15 subjects each; for each fold we performed a multinomial logistic regression with the first three gradients as predictor variables and networks as outcome variables. Beta values derived from the training set were used to predict probabilities of each network in the testing set. Each vertex was assigned to the network with the highest probability. Additionally, we derived beta values from the entire *HCP-Discovery* dataset and used these to predict functional networks from *HCP-Replication* and *MICs* gradients.

To further assess the relationship between structural gradients and edgewise functional connectivity, we used a decision tree with binary splits for regression. Similar to the network prediction, training and testing was performed both with 5-fold cross validation as well as training on HCP-Discovery and testing on the other datasets. Model training was performed with the *fitrtree* function as implemented in MATLAB R2019b with a minimum leaf size of 20, a maximum number of splits of 20, and otherwise default parameters.

### Evolutionary Analyses

We tested for associations between our gradients and two markers of evolutionary change between humans and macaques: functional homology and areal expansion. Both measures were presented in a prior paper (Xu et al., 2020), hence we only provide a short overview here. Functional homology is a measure for the functional similarity of a human brain area with its macaque counterpart. It is computed based on the maximum cosine similarity of functional gradient profiles within a 12mm search light around the corresponding human/macaque vertices. An areal expansion map shows the relative expansion of human cortex compared to macaques. It is computed by dividing the local area of human cortex by the corresponding area of macaque cortex where correspondence was defined based on functional homology. We tested for associations between these two markers and the structural gradients using Moran spectral randomization (Wagner & Dray, 2015).

## Acknowledgements

Reinder Vos de Wael was funded by studentships from the Savoy foundation for Epilepsy and the Richard and Ann Sievers award. Dr. Jessica Royer was supported by a fellowship from the Canadian Open Neuroscience Platform (CONP) and CIHR. Shahin Tavakol was funded by a studentship from McGill University’s Faculty of Medicine. Dr Casey Paquola was funded through a postdoctoral fellowship of the Fonds de la Recherche du Quebec - Santé (FRQ-S). Dr Oualid Benkarim was funded by a Healthy Brains for Healthy Lives (HBHL) postdoctoral fellowship. Sara Larivière was funded by the Canadian Institutes of Health Research (CIHR). Dr Jonathan Smallwood was supported by the European Research Council (WANDERINGMINDS-ERC646927). Dr Boris Bernhardt acknowledges research support from the National Science and Engineering Research Council of Canada (NSERC Discovery-1304413), the Canadian Institutes of Health Research (CIHR FDN-154298), SickKids Foundation (NI17-039), Azrieli Center for Autism Research (ACAR-TACC), and the Tier-2 Canada Research Chairs program.

Data were provided, in part, by the Human Connectome Project, WU-Minn Consortium (Principal Investigators: David Van Essen and Kamil Ugurbil; 1U54MH091657) funded by the 16 NIH Institutes and Centers that support the NIH Blueprint for Neuroscience Research; and by the McDonnell Center for Systems Neuroscience at Washington University.

