## Supplementary Figures for "Microstructural underpinnings and macroscale functional implications of temporal lobe connectivity gradients"

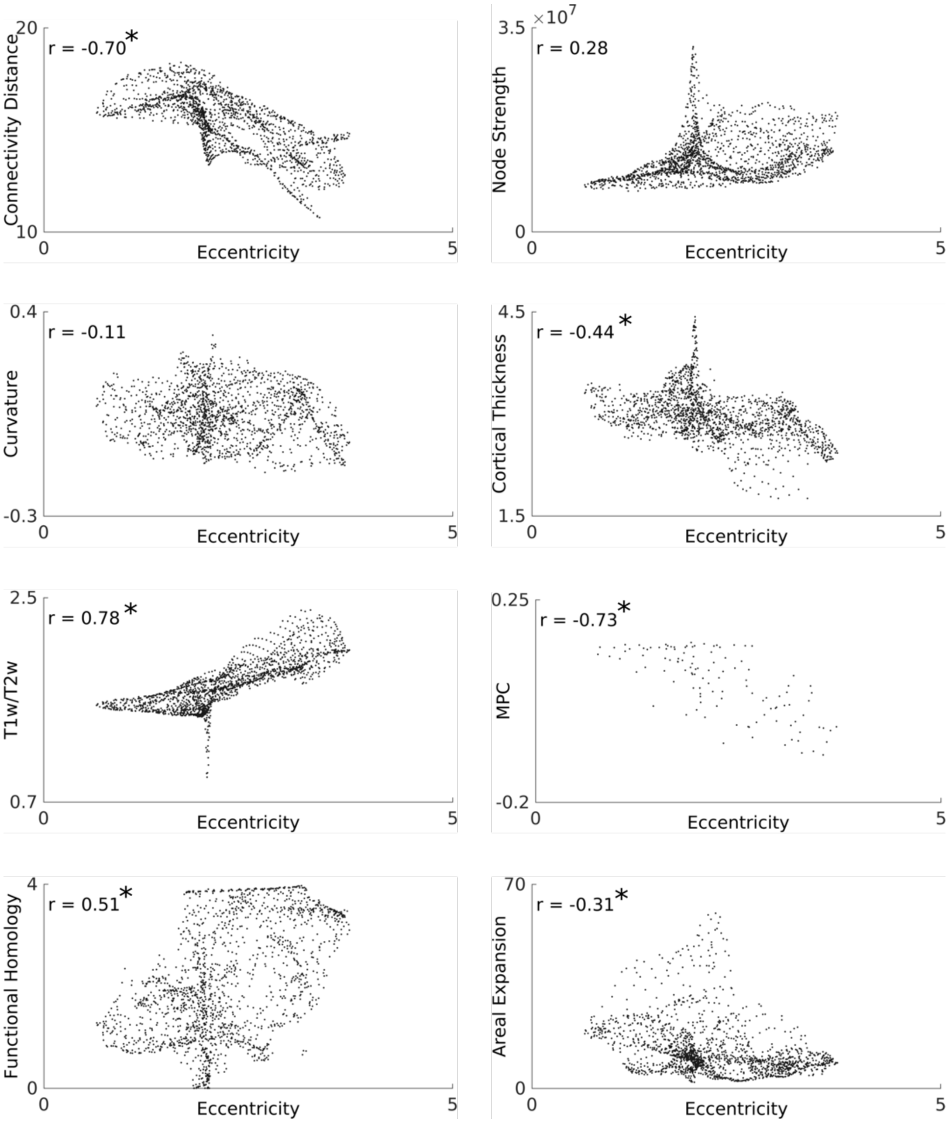


**Supplementary Fig. 1.** Main right hemispheric results of *HCP-Discovery*. Stars denote significant results ($p_{moran}$<0.05).

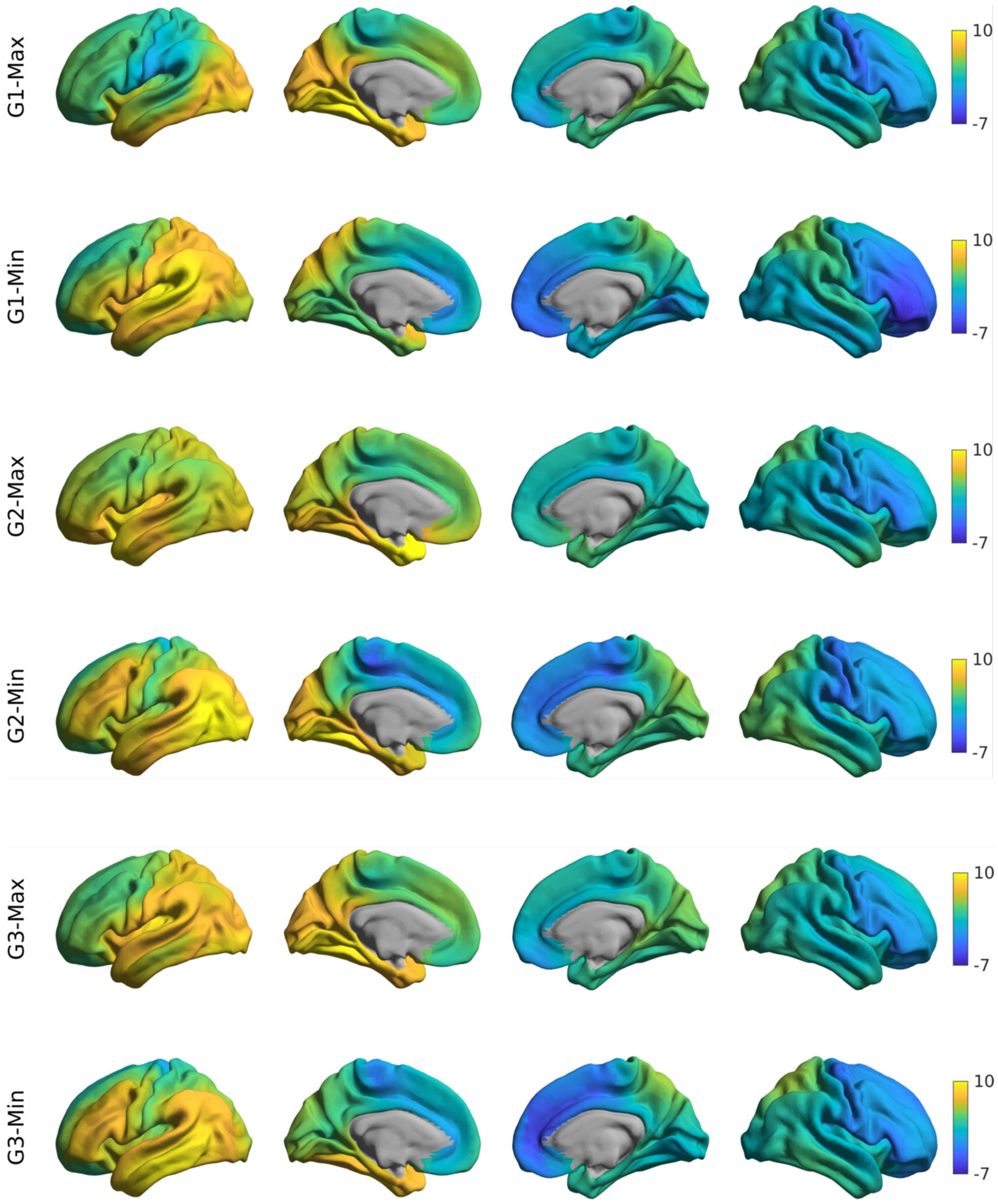


**Supplementary Fig. 2.** Connectivity profiles of the top (G-max) bottom (G-min) 10% of the left-hemispheric gradients. Data were log-transformed for visualization purposes only.

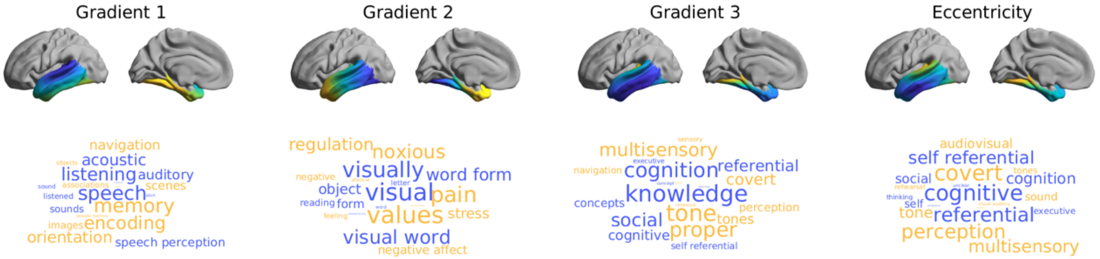


**Supplementary Fig 3.** Associations between gradients/eccentricity and meta-analytic cognitive terms derived from Neurosynth. Terms in blue are associated with the lower (blue) end of the gradients and yellow terms with the higher (yellow end).
